# Change in protein, oil and fatty acid contents in soybeans (*Glycine max* (L.) Merr.) of different seed coat colors and seed weight

**DOI:** 10.1101/2021.02.10.430590

**Authors:** Yu-Mi Choi, Hyemyeong Yoon, Myoung-Jae Shin, Yoonjung Lee, On Sook Hur, Bong Choon Lee, Bo-Keun Ha, XiaoHan Wang, Kebede Taye Desta

**Affiliations:** National Agrobiodiversity Center, National Institute of Agricultural Sciences, Rural Development Administration, Jeonju 54874, Korea; Crop Foundation Division, National Institute of Crop Science, Wanju 55365, Korea; Division of Plant Biotechnology, Chonnam National University, Gwangju 61186, Korea; Department of Applied Chemistry, Adama Science and Technology University, Adama 1888, Ethiopia

**Author notes:** Corresponding author (KTD).

## Abstract

Soybean seeds are one of the best sources of plant-based high-quality proteins and oils. The contents of these metabolites are affected by both environmental and genetic factors. In this study, 49 soybean germplasms were cultivated in Korea, the contents of total protein, total oil and five fatty acids were determined, and the influences of seed coat color and seed weight on each were assessed. The total protein and total oil contents were evaluated using Kjeldahl and Soxhlet methods and were in the ranges of 36.28-44.19% and 13.45-19.20%, respectively. Moreover, the contents of individual fatty acids were determined as area percentage from acquired gas-chromatography peaks. The contents of palmitic, stearic, oleic, linoleic and linolenic acids were in the ranges of 9.90-12.55, 2.45-4.00, 14.97-38.74, 43.22-60.26, and 5.37-12.33%, respectively and each significantly varied between the soybean germplasms. Unlike total oil and fatty acid contents, total protein content was not significantly affected by both seed coat color and seed weight. Cluster analysis grouped the soybeans into two classes with notable content differences. Fatty acids were the main factors for the variabilities seen between the soybean germplasms as observed in the principal component analysis. Correlation analysis revealed a significant but negative association between total oil and total protein contents (*r* = -0.714, *p* < 0.0001). Besides, a trade-off relationship was observed between oleic acid and linoleic acid (*r* = -0.936, *p* < 0.0001) which was reflected with respect to both seed coat color and seed weight. Among all colored soybeans, pale-yellow soybeans had the highest and the lowest levels of oleic acid and linoleic acid, respectively each being significantly different from the rest of colored soybeans (*p* < 0.05). Likewise, oleic acid content increased with seed weight while that of linoleic acid decreased with seed weight (*p* < 0.05). In general, this study showed the significance of seed coat color and seed weight to discriminate soybean genotypes, mainly in terms of their fatty acid contents. Moreover, the soybean germplasms with distinct characters and fatty acid contents identified in this study could be important genetic resources for cultivar development.

## Introduction

Soybean (*Glycine max* (L.) Merr.) has been extensively cultivated in many parts of the world. Currently, Brazil is the leading soybean producer worldwide followed by the United States and Argentina (http://www.fao.org/faostat).

Soybean seeds are considered as one of the best sources of plant-based high-quality proteins and oils. Other health-promoting secondary metabolites such as phenolic acids, isoflavones, and anthocyanins are also ubiquitous in soybean [1, 2]. Several studies specified that the protein and oil contents in soybean seeds account for 35-55 % and 13-26% on a dry seed weight basis, respectively [3, 4]. It is widely documented that soybean proteins contain all the essential amino acids. Owing to these, soybean protein has been used in the human diet in a variety of forms such as infant formula, protein isolates and flours [5, 6]. Likewise, soybean oil contains five prominent fatty acids including two saturated fatty acids (palmitic acid (16:0) and stearic acid (18:0)) and three unsaturated fatty acids (oleic acid (18:1), linoleic acid (18:2), and linolenic acid (18:3)). The high level of unsaturated fatty acids, mainly of linoleic and oleic acids, denotes the role of soybean in preventing many ailments including cancer, diabetes, and cardiovascular diseases [7, 8]. Moreover, soybean oil is becoming a major feedstock for biodiesel production. In 2019 alone, 77% of the total biodiesel production was derived from vegetable oils and out of that soybean oil contributed 30% [9]. Researchers have also been modifying the levels of targeted fatty acids to enhance the performance of soybean oil in biofuel blends [10]. In general, the protein, oil and fatty acids contents in soybean are among the main driving factors for its increasing use in food, pharmaceutical, and biofuel industries. To meet the growing demands in various sectors, developing soybean genotypes with improved or altered protein, oil and fatty acid contents has become an important breeding objective [11, 12].

The contents of soybean metabolites, in general, are affected by both environmental and genetic factors. Temperature, growing location, year of cultivation, farming condition and solar radiation are among the major environmental factors and much work has been conducted on the connection between these factors and soybean metabolites [5, 13, 14]. From a genetic standpoint, seed-related characters such as seed coat color and seed weight are the main traits that determine soybean seed quality. Researchers identified several minor and major genes that regulate seed coat color and seed weight in soybean which influence not only the yield but also the metabolite contents in matured seeds [2, 15, 16]. Moreover, previous studies documented that seed coat color and seed weight influence the levels of other secondary metabolites including isoflavones, anthocyanins, tocopherols and phenolic compounds in soybeans [13, 17-19]. Unlike these, however, studies that investigate the effects of seed-related characters on protein, oil and fatty acid contents in soybeans are still limited [3, 8, 20, 21]. In general, analyzing the effects of these factors on protein, oil, and fatty acid contents could be a significant input for the development of improved soybean cultivars [16]. Besides, such studies are useful to identify desirable soybean characteristics that lift consumers’ preferences [22].

As in other countries, there is growing attention towards developing improved soybean cultivars in Korea. Previously, several researchers assessed the levels of metabolites in soybean germplasms cultivated in Korea and also attempted to evaluate the effects of some environmental and genetic factors although such studies are infrequent [3, 8, 13, 23]. This study intended to assess the influences of seed coat color and seed weight on the contents of total protein, total oil, and five fatty acids (palmitic acid, stearic acid, oleic acid, linoleic acid and linolenic acid) in 49 soybean germplasms recently cultivated in Korea.

## Materials and methods

### Plant materials

The seeds of 49 soybean germplasms were obtained from the gene bank of the National Agrobiodiversity Center, Rural Development Administration (www.genebank.rda.go.kr, Jeonju, Republic of Korea). The soybeans were cultivated between June and November of 2019 under similar environmental conditions in an experimental field located at the center [24]. Matured seeds were hand-harvested and grouped as black (*n*=24), yellow (*n*=10), yellowish-green (*n*=5), green with black spot (*n*=4), pale-yellow (*n*=4), and green (*n*=2) based on their seed coat color (**Fig 1**). Besides, the soybeans were grouped as small (<13 g; *n*=8), medium (13-24 g; *n*=14), and large (>24 g; *n*=27) based on their one-hundred seeds weight (HSW) [23]. Samples of whole seed soybeans from each class were dried in Bionex Convection oven (Vision Scientific, Daejeon, Korea) for three days at 50 °C. The dried seed samples were then powdered, passed through a 315 *μ*m sieve, and used for protein, oil and fatty acid analysis. Powdered seed samples were stored at -20 °C when not used. Additional information related to name, introduction (IT) number, seed coat color, seed weight and sample code of each soybean germplasm can be found in S1 Table.

**Fig 1.**
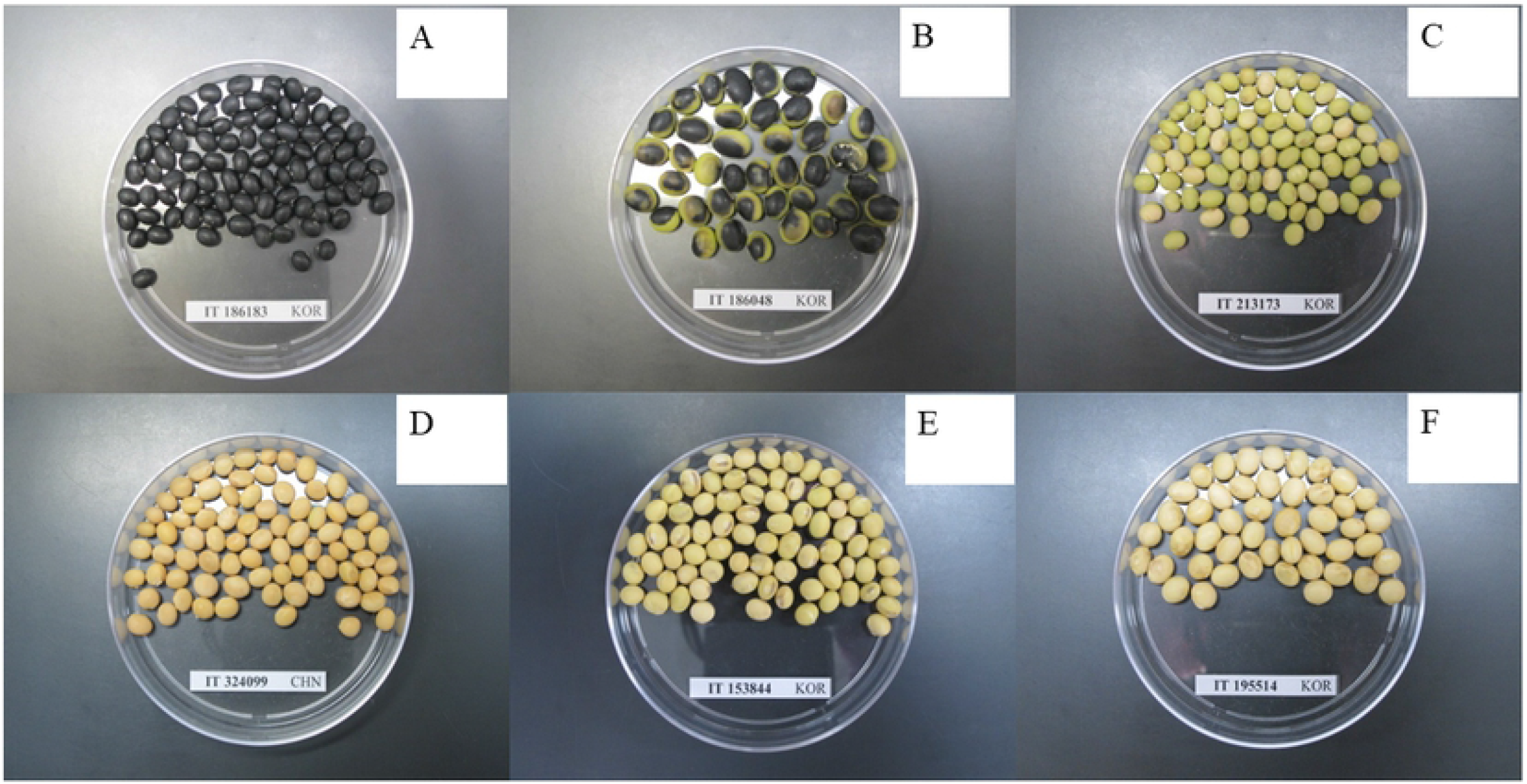
Seed samples of soybean germplasms of different seed coat colors. A: Black, B: Green with black spot, C; Green, D: Yellow, E: Yellowish-green, F: Pale-yellow

### Chemicals and reagents

All the chemicals and reagents used in this study were of analytical grade and used without further purification steps. Water and *n*-hexane were purchased from Thomas Scientific (Philadelphia, PA, USA) and Fisher Scientific (Pittsburgh, PA, USA), respectively. All the other chemicals and reagents including methanol, anhydrous sodium sulfate (Na^2^SO^4^), sulfuric acid (H^2^SO^4^), boron trifluoride-methanol (BF^3^-MeOH) solution, sodium hydroxide (NaOH) and standards of palmitic acid, stearic acid, oleic acid, linoleic acid and linolenic acid were obtained from Sigma Aldrich (St. Louis, MO, USA).

### Determination of total protein content

The protein content in seeds of each soybean was determined according to the Kjeldahl method using a Kjeltec instrument equipped with an *auto*-digester (FOSS, Tecator, Hoganas, Sweden) [3]. In brief, 0.5 g of powdered whole seed sample was put into a digestion tube. The sample was digested using 12 mL of concentrated H^2^SO^4^ and two pellets of selenium catalyst for 1 h. The digestion tube was then removed, cooled to room temperature (∼25°C), and processed using an automatic Kjeltec analyzer (FOSS, Tecator, Hoganas, Sweden) which was capable of distillation and colorimetric titration. The total protein content (%) was *auto*-computed by multiplying the nitrogen content by a factor of 6.25 (the standard Kjeldahl factor).

### Determination of total oil content

The total oil content was determined by a standard Soxhlet extraction procedure using a Soxtec extraction system (FOSS, Tecator, Hoganas, Sweden) [25]. In brief, 0.7 g of powdered soybean whole seed sample was placed in an extraction thimble and mixed with 50 mL of *n*-hexane. The thimble containing the mixture was loaded in an extraction unit maintained at 135°C and the boiling, rinsing and recovery times were automated at 30, 60, and 20 min, respectively. After extraction, the oil was cooled to room temperature in a desiccator. The percentage of total oil content was determined as the mass ratio of obtained oil and extracted seed sample on a dry weight basis from triplicate repetitions.

### Methylation of fatty acids and GC-FID analysis

Before analysis, the fatty acids were esterified to the corresponding fatty acid methyl esters (FAMEs) through transmethylation [26]. Specifically, 50 *µ*L of the extracted oil was placed in a 50 mL capacity conical tube and 2 mL of 0.5M NaOH solution in methanol was added. The mixture was vortex-mixed and heated in a water bath at 80°C for 10 min. Then, the mixture was cooled to room temperature and the reaction cycle was repeated one more time after the addition of 2 mL of BF^3^-MeOH solution. In the final step, 7 mL of *n*-hexane and 2 mL of water were sequentially added, the mixture was vortexed, and centrifuged (4°C, 3000 rpm, for 10 min). The upper organic layer was retained, dried over anhydrous Na^2^SO^4^, and stored in a vial. For analysis, 10 *µ*L of FAMEs sample was injected into QP2010 gas chromatography (GC) (Shimadzu, Kyoto, Japan) equipped with a flame ionization detector (FID). The separation of FAMEs was achieved using an HP-INNOWAX column (30 m x 0.250 mm x 0.25 *µ*m) and the temperature gradient was optimized to start with 100°C followed by an increase to 240°C at a rate of 6.5°C/min for a total of 25 min. The temperatures of the injection port and the detector were each set at 250°C. Helium was used as a carrier gas at a flow rate of 1.5 mL/min and a split ratio of 50:1. Individual FAMEs were identified by comparing the retention times of the corresponding standards and quantified as the percentage of the total fatty acid from peak areas of the acquired GC-chromatograms.

### Data analysis

In this study, all measurements were made in triplicates and results are expressed as mean ± SD. One-way analysis of variance was computed using xlstat-software (Addinsoft, Long Island, NY, USA) and differences were considered statistically significant at a probability value of < 0.05 (*p* < 0.05). Boxplots, cluster analysis, principal component analysis (PCA) and correlation analysis were performed using R-software *ver*. 4.0.2 (https://www.r-project.org/).

## Results and discussion

### Metabolite contents in the soybean germplasms

The contents of each metabolite for all the soybeans are presented in S1 Table and significant variations were observed between the soybean germplasms. With a mean of 39.55%, the total protein content ranged between 36.28% in soybean YG49 and 44.19% in soybean YG48 (*p* < 0.05). The total oil content was in the range of 13.45-19.20% with a mean of 11.03%. Soybeans BL5 and YG48 had the highest and the lowest total oil contents, respectively (*p* < 0.05). These findings were in agreement with the results reported by Cho et al. [23] who found a total protein content in the range of 38.97-44.46% and a total oil content in the range of 13.45-20.38% in soybeans grown in Korea. In another study, Lee et al. [8] reported a total protein content that ranged between 46.01 and 48.44% which was less wide but higher compared to our findings and a total oil content that ranged between 15.74 and 17.20% which was less wide than our observation. Moreover, Kumar et al. [22] found a total protein content that ranged between32.20 and 42.10% and a total oil content that ranged between 15.40 and 22.00% in Indian soybeans, whereas Qin et al. [27] found total protein and total oil contents in the ranges of 31.79-49.78 and 14.17-22.76% with means of 39.32 and 20.15%, respectively in Chinese soybeans. In addition to the difference in genotypes, variations in farming conditions, temperature and year of cultivation could be the factors that cause the observed inconsistencies [21, 28].

The contents of the five individual fatty acids, total saturated fatty acid (TSFA) and total unsaturated fatty acid (TUFA) also significantly varied among the soybeans (S1 Table). The contents of palmitic, stearic, oleic, linoleic and linolenic acids were in the ranges of 9.90-12.55, 2.45-4.00, 14.97-38.74, 43.22-60.26, and 5.37-12.33% with means of 11.03, 3.15, 22.74, 54.75,and 8.33%, respectively. The ranges of the unsaturated fatty acids found in this study were slightly wider than those reported by Shin et al. [29] who previously studied 18 soybean cultivars grown in Korea. It is evident from S1 Table that the level of linoleic acid was dominant in every soybean followed by oleic acid, which agreed with previous findings [8, 30, 31]. Oleic acid content exhibited the highest coefficient of variation (CV) (19.59%) demonstrating a high variability between the soybeans, whereas linoleic acid content displayed the lowest CV (6.04%). Specific to saturated fatty acids, the content of palmitic acid was dominant over stearic acid as observed in other studies [29, 30]. Among all the soybeans, soybeans BL19 and BL17 had the highest, whereas soybeans YG47 and PY33 had the lowest palmitic and stearic acid contents, respectively. Soybean PY31 had the highest oleic acid and the lowest linoleic acid contents, whereas soybean BL16 had the lowest oleic acid and the highest linoleic acid contents at the same time indicating the trade-off relationship between these two fatty acids [8, 29, 32]. Soybean PY31 also displayed the lowest level of linolenic acid while soybean BL19 had the highest level. Linolenic acid, a polyunsaturated fatty acid, is characterized by oxidative instability causing rancidity and reduced shelf-life of soybean oil and soy-products. Due to these, attempts have been made to reduce the level of linolenic acid in soybeans through breeding. Together, increasing the level of oleic acid is usually taken into account to attain the best breeds [33-35]. In this study, soybeans BL3, BL12, GR30, PY31, PY33, and PY34 were typically found to contain a low level of linolenic acid (< 7.00%) and a high level of oleic acid (> 25%) signifying their potential for the production of good quality soybean oil. Besides, they possibly provide a wide spectrum of options if considered during breeding of improved soybean cultivars.

### Metabolite content variations with respect to seed coat color

To view the influences of seed color, the soybeans were grouped based on their seed coat color as described before. The box plots in Fig 2 show the variation of each metabolite content with respect to seed coat color and the corresponding numerical value can be read from Table 1. The average protein content was in the order of green (40.18%) > black (39.97%) > yellow (39.62%) > yellowish-green (39.04%) > pale-yellow (39.04%) > green with black spot (37.99%) soybeans. Likewise, the average oil content decreased in the order of green with black spot (18.04%) > pale-yellow (17.59%) > yellowish-green (16.97%) > green (16.81%) > yellow (16.55%) > black (16.43%) soybeans. The average protein and oil contents found in black, yellow, and green soybeans were slightly lower than those reported by Cho et al. [3] which could be attributed to the differences in genotypes and growing conditions. Besides, there was an insignificant difference in total protein content between the six colored soybeans (Fig 2). With regard to total oil content, only black and green with black spot soybeans significantly varied with each other (*p* < 0.05). Previous studies also did not notice significant variations of total protein and oil contents between other colored soybeans grown in Korea [3, 8]. In another study, Zarkadas et al. [36] noticed small but significant differences in protein content between yellow and brown soybeans cultivars adapted in Canada. Moreover, Redondo-Cuenca et al. [37] found significant variations in protein and fat contents between ecological and transgenic yellow and green soybeans of different countries of origin. In general, our findings signify that soybean genotypes should still be evaluated individually, not based on the color of their seed coats, for their use in soy-protein formulations and dietary oil production.

**Table 1.**

| **Metabolite** | **Values (%)** | **Seed coat color** |  |  |  |  |  | **Seed weight** |  |  |
| --- | --- | --- | --- | --- | --- | --- | --- | --- | --- | --- |
|  |  | **Black**<br>**(n=24)** | **GBS**<br>**(n=4)** | **Green**<br>**(n=2)** | **PY**<br>**(n=4)** | **YG**<br>**(n=5)** | **Yellow**<br>**(n=10)** | **Small**<br>**(n=8)** | **Medium**<br>**(n=14)** | **Large**<br>**(n=27)** |
| **Total protein** | Minimum | 36.53 | 36.82 | 37.68 | 37.60 | 36.28 | 36.46 | 38.84 | 37.60 | 36.28 |
|  | Maximum | 42.65 | 38.62 | 42.68 | 40.32 | 44.19 | 41.63 | 42.14 | 42.68 | 44.19 |
|  | Mean | 39.97 | 37.99 | 40.18 | 38.79 | 39.04 | 39.62 | 40.62 | 39.31 | 39.36 |
|  | CV | 3.86 | 2.20 | 8.79 | 3.04 | 7.74 | 3.80 | 2.83 | 4.01 | 4.89 |
| **Total oil** | Minimum | 14.16 | 17.26 | 15.80 | 16.37 | 13.45 | 15.17 | 14.35 | 15.80 | 13.45 |
|  | Maximum | 19.20 | 18.82 | 17.83 | 18.46 | 18.98 | 18.06 | 15.99 | 18.98 | 19.20 |
|  | Mean | 16.43 | 18.04 | 16.81 | 17.59 | 16.97 | 16.55 | 15.16 | 17.32 | 16.93 |
|  | CV | 8.09 | 3.55 | 8.53 | 5.05 | 14.38 | 5.66 | 3.20 | 5.79 | 8.19 |
| **Palmitic acid** | Minimum | 9.98 | 10.11 | 11.49 | 9.94 | 9.90 | 10.54 | 10.54 | 9.90 | 9.94 |
|  | Maximum | 12.55 | 10.95 | 12.06 | 11.49 | 11.19 | 12.19 | 12.26 | 11.95 | 12.55 |
|  | Mean | 11.10 | 10.61 | 11.77 | 10.84 | 10.55 | 11.17 | 11.33 | 10.81 | 11.05 |
|  | CV | 6.67 | 3.43 | 3.44 | 6.05 | 5.29 | 5.09 | 4.38 | 6.47 | 6.29 |
| **Stearic acid** | Minimum | 2.55 | 3.19 | 2.64 | 2.45 | 2.82 | 2.80 | 2.55 | 2.87 | 2.45 |
|  | Maximum | 4.00 | 3.41 | 3.70 | 3.15 | 3.47 | 3.60 | 3.89 | 4.00 | 3.45 |
|  | Mean | 3.20 | 3.28 | 3.17 | 2.72 | 3.11 | 3.14 | 3.42 | 3.26 | 3.01 |
|  | CV | 11.36 | 2.89 | 23.69 | 11.14 | 8.09 | 7.67 | 13.55 | 9.35 | 8.17 |
| **Oleic acid** | Minimum | 14.97 | 18.76 | 16.86 | 25.33 | 18.11 | 16.03 | 14.97 | 16.02 | 18.27 |
|  | Maximum | 28.67 | 23.82 | 28.68 | 38.74 | 28.22 | 28.33 | 21.00 | 28.67 | 38.74 |
|  | Mean | 21.18 | 21.36 | 22.77 | 30.70 | 23.58 | 23.42 | 18.88 | 22.73 | 23.89 |
|  | CV | 13.10 | 12.25 | 36.70 | 20.43 | 15.74 | 19.33 | 9.77 | 18.07 | 19.33 |
| **Linoleic acid** | Minimum | 51.99 | 54.43 | 49.77 | 43.22 | 51.43 | 49.22 | 54.14 | 50.71 | 43.22 |
|  | Maximum | 60.26 | 59.02 | 58.10 | 53.86 | 59.15 | 59.05 | 60.26 | 59.05 | 59.02 |
|  | Mean | 55.83 | 56.54 | 53.94 | 49.30 | 54.63 | 53.85 | 57.44 | 55.34 | 53.65 |
|  | CV | 3.61 | 3.85 | 10.92 | 9.73 | 5.91 | 6.25 | 3.28 | 4.87 | 6.44 |
| **Linolenic acid** | Minimum | 5.77 | 7.53 | 6.85 | 5.37 | 7.00 | 6.90 | 8.28 | 5.77 | 5.37 |
|  | Maximum | 12.33 | 9.04 | 9.85 | 8.15 | 9.63 | 10.35 | 10.04 | 10.35 | 12.33 |
|  | Mean | 8.68 | 8.21 | 8.35 | 6.44 | 8.14 | 8.42 | 8.93 | 7.86 | 8.40 |
|  | CV | 15.95 | 9.47 | 25.37 | 18.61 | 13.07 | 12.87 | 7.53 | 17.84 | 16.91 |
| **TSFA** | Minimum | 13.52 | 13.52 | 14.70 | 12.67 | 12.99 | 13.45 | 14.26 | 13.06 | 12.67 |
|  | Maximum | 15.55 | 14.21 | 15.19 | 14.63 | 14.31 | 15.17 | 15.34 | 15.19 | 15.55 |
|  | Mean | 14.31 | 13.89 | 14.95 | 13.57 | 13.66 | 14.31 | 14.75 | 14.07 | 14.06 |
|  | CV | 4.90 | 2.16 | 2.32 | 5.95 | 4.47 | 4.50 | 2.60 | 5.18 | 4.97 |
| **TUFA** | Minimum | 84.45 | 85.79 | 84.81 | 85.37 | 85.69 | 84.83 | 84.66 | 84.81 | 84.45 |
|  | Maximum | 86.94 | 86.48 | 85.30 | 87.33 | 87.01 | 86.55 | 85.74 | 86.94 | 87.33 |
|  | Mean | 85.69 | 86.11 | 85.05 | 86.43 | 86.34 | 85.69 | 85.25 | 85.93 | 85.94 |
|  | CV | 0.82 | 0.35 | 0.41 | 0.93 | 0.71 | 0.75 | 0.45 | 0.85 | 0.81 |

Variation of total protein, total oil and fatty acid contents in soybeans with respect to seed coat color and seed weight. GBS: Green with black spot, PY: Pale-yellow; TSFA: Total saturated fatty acid; TUFA; Total unsaturated fatty acid; YG: Yellowish- green.

**Fig 2.**
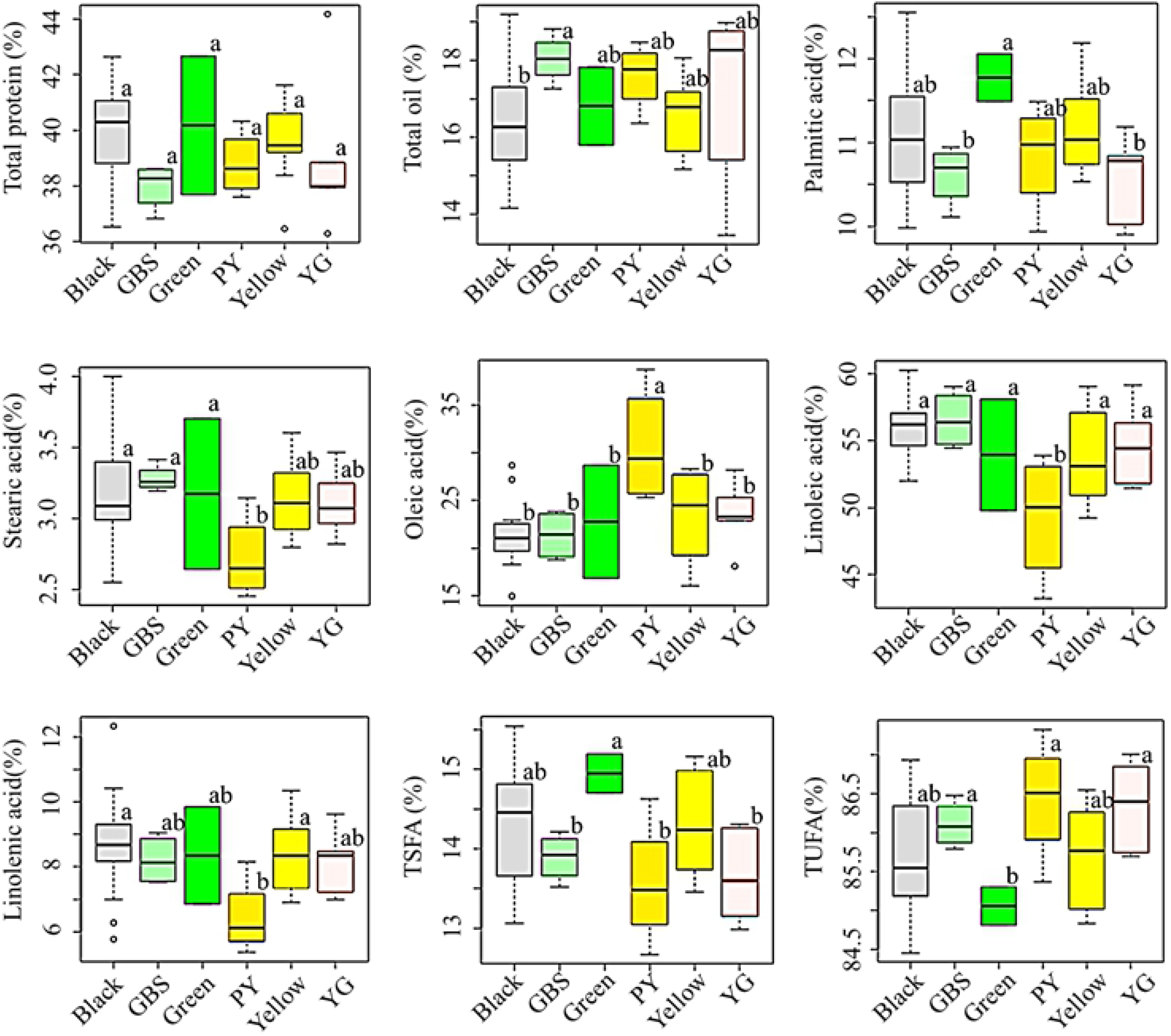
Variation of total protein, total oil and individual fatty acid contents in soybeans according to seed coat color. GBS: Green with black spot, PY: Pale-yellow; TSFA: Total saturated fatty acid; TUFA; Total unsaturated fatty acid; YG: Yellowish-green. Different letters on box plots in a category represent means that are significantly different (*p* < 0.05).

Fig 2 again shows the changes in individual fatty acid contents among the six colored soybean groups. The levels of palmitic acid and stearic acid were the highest in green (11.77%) and green with black spot (3.28%) soybeans, and the lowest in yellowish-green (10.55%) and pale-yellow soybeans (2.72%), respectively (Table 1). The palmitic acid content in green with black spot and yellowish-green soybeans was significantly different from that of green soybeans, whereas stearic acid content in pale-yellow soybeans was significantly different from black, green and green with black spot soybeans (*p* < 0.05). With regard to unsaturated fatty acids, pale-yellow soybeans had the highest level of oleic acid (30.70%) and the lowest level of linoleic acid (49.30%) each being significantly different from the rest colored soybeans (*p* < 0.05). These observations further attest to the contrasting relationship between oleic acid and linoleic acid [27]. Opposite to pale-yellow soybeans, black soybeans had the lowest oleic acid content (21.18%) while green with black spot soybeans had the highest linoleic acid content (56.54%). The black and yellow soybeans studied in this study had a lower average oleic acid content but a higher average linoleic acid contest than soybeans of similar seed coat colors reported by Shin et al. [29]. The average linolenic acid content was the highest in black soybeans (8.68%) and the lowest in pale-yellow soybeans (6.44%). The level of linolenic acid in pale-yellow soybeans was significantly different from that of black and yellow soybeans (*p* < 0.05). In other studies, variable results were reported with respect to fatty acid contents between colored soybeans although such studies are still limited. For instance, Cho et al. [3] failed to note significant variations in the contents of all five fatty acids between black, brown, green and yellow soybeans. In a later study, however, significant variations were observed in the levels of all fatty acids between these colored soybeans [8]. In general, our results suggest that seed coat color could be an important agronomic character to categorize soybeans with respect to the levels of fatty acids. Moreover, the pale-yellow soybeans identified in this study could be important genetic resources owing to their simultaneous low average linolenic acid and high average oleic acid contents.

### Metabolite content variations with respect to seed weight

Seed weight, determined in-terms of HSW, is an important agronomical character, and genes associated with it have been targeted to improve the quality of soybean seeds [15]. Previously, few studies investigated the influence of seed weight on protein, oil, and fatty acid contents and inconsistent findings were reported [8, 38,39]. In this study, the soybeans were grouped as small, medium and large seeds based on their HSW and the influence of seed weight on the content of each metabolite was assessed (Fig 3, Table 1). The average total protein content decreased in the order of small (40.62%) > large (39.63%) > medium (39.31%). However, no significant difference was observed between any of the three groups. The total oil content decreased in the order of medium (17.32%) > large (16.93%) > small (15.16%) soybeans the latter being significantly different from the other two groups (*p* < 0.05).

**Fig 3.**
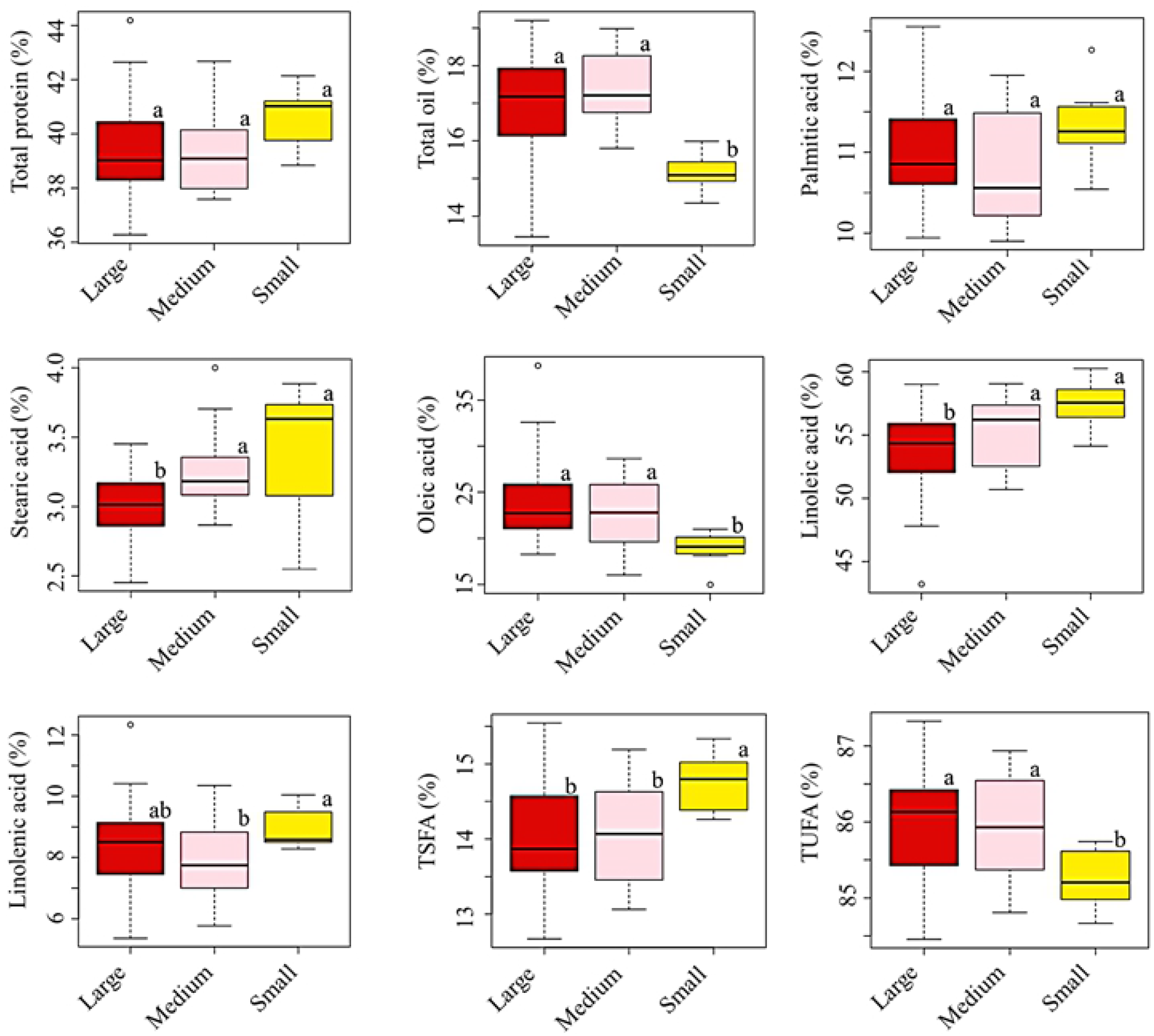
Variation of total protein, total oil and fatty acid contents in soybeans with respect to seed weight. TSFA: Total saturated fatty acid; TUFA; Total unsaturated fatty acid. Different letters on box plots in a category represent means that are significantly different (*p* < 0.05).

With respect to individual fatty acids, the contents of palmitic acid and linolenic acid were each the highest in small seeds (11.33 and 8.93%, respectively) followed by large (11.05 and 8.40%, respectively) and medium (10.81 and 7.86%, respectively) seeds. There was an insignificant difference in palmitic acid content between small, medium and large seeds (Fig 3). Stearic acid and linoleic acid contents decreased with seed weight each being in the order of small > medium > large seeds. The levels of these fatty acids in large seeds were significantly different from the rest two groups (*p* < 0.05). Opposite to this, the level of oleic acid increased with seed weight and large seeds had the highest level (23.89%) followed by medium (22.73%) and small seeds (18.88%) (*p* < 0.05). Moreover, the variations of TSFA and TUFA between small, medium and large seeds were significant (*p* < 0.05). The overall observation was consistent with Lee et al. [8] who also noted significant variations in the levels of all fatty acids with respect to seed weight. Soybeans of a specific seed coat color were also grouped and analyzed based on their HSW (S2 Table). In black soybeans, significant variations were observed in all but total protein and linoleic acid contents. Moreover, the variations of oleic acid and linoleic acid contents between small, medium and large seeds of pale-yellow soybeans were significant (*p* < 0.05). In general, our results suggest that seed weight could be an important agronomical parameter to distinguish soybean genotypes with respect to their fatty acid contents.

### Cluster, principal component and correlation analysis

To view the distribution and relationship of the soybeans over the analyzed metabolites, a combined heatmap and cluster analysis, PCA and correlation analysis were conducted using the whole data set. It is evident from Fig 4A that the soybeans were grouped into two main parts with notable metabolite content differences. The first group was characterized by low levels of oleic acid and total oil, whereas the second group was characterized by low levels of palmitic acid and TSFA. Besides, the majority of the soybeans in the first group had high levels of palmitic acid and TSFA while those in the second cluster had high levels of oleic acid, TUFA, and total oil. Interestingly, all small soybeans and the majority of black soybeans were clustered in the first group, whereas all pale-yellow soybeans, green with black spot soybeans and the majority of large soybeans were clustered in the second group. Moreover, the heatmap confirmed the negative relationship between oleic acid and linoleic acid irrespective of seed coat color and seed weight. The distribution of the soybeans was further viewed by PCA which yielded three components with eigenvalues > 1 (Table 2). The first two components (PC1 and PC2) which explained 72.23% of the cumulative variance were taken for analysis. The two most notable observations in the PCA were the clustering of pale-yellow soybeans (Fig 4B) and small soybeans (Fig 4C) which were in accordance with the heatmap results. The corresponding loading plot revealed oleic acid and palmitic acid as the most discriminative variables along PC1 and PC2, respectively (Fig 4D, Table 2). These observations signify that oleic acid and palmitic acid could selectively be targeted to categorize a large population of soybeans [32].

**Table 2.**
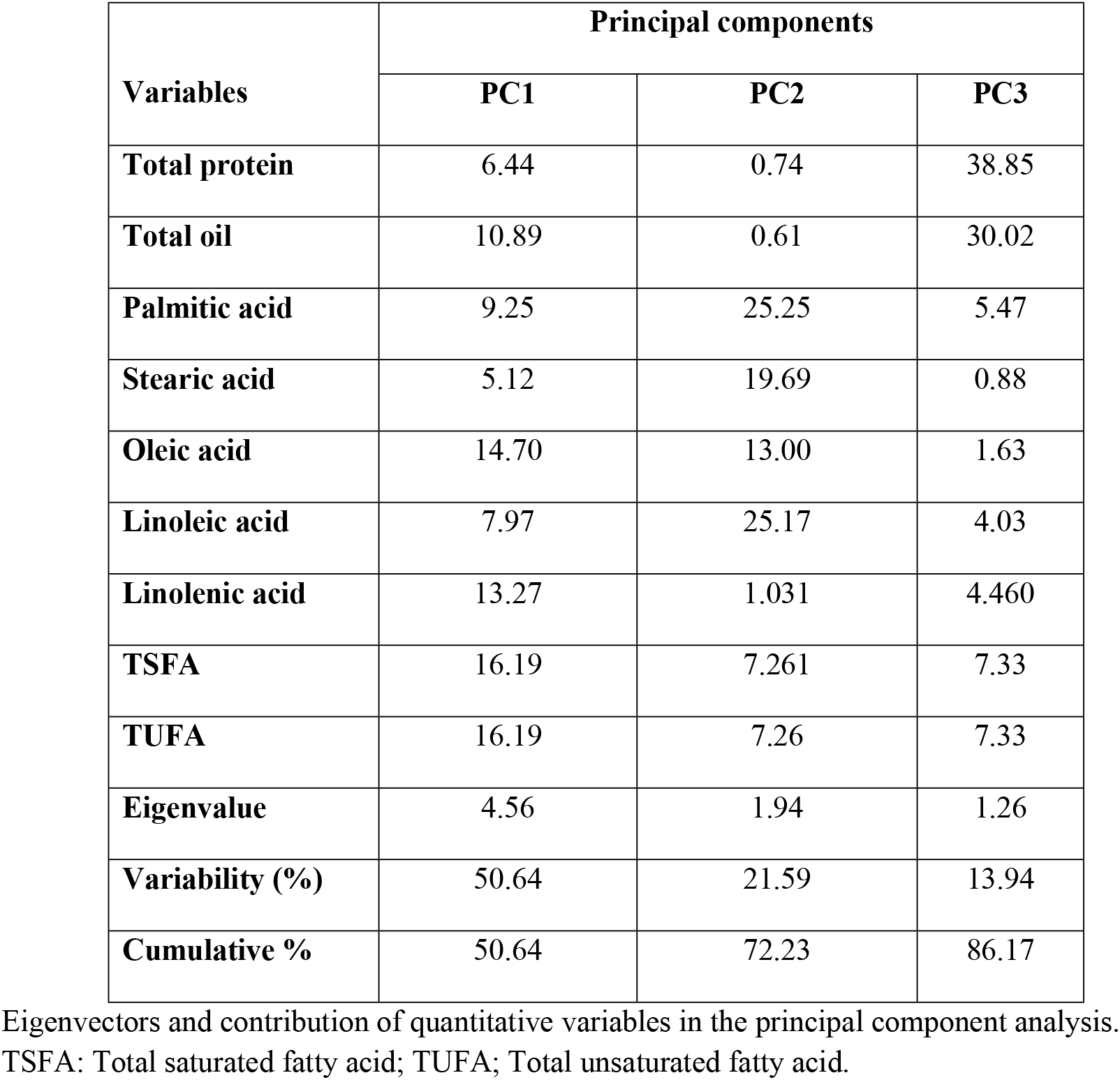

| Variables | Principal components |  |  |
| --- | --- | --- | --- |
|  | PC1 | PC2 | PC3 |
| Total protein | 6.44 | 0.74 | 38.85 |
| Total oil | 10.89 | 0.61 | 30.02 |
| Palmitic acid | 9.25 | 25.25 | 5.47 |
| Stearic acid | 5.12 | 19.69 | 0.88 |
| Oleic acid | 14.70 | 13.00 | 1.63 |
| Linoleic acid | 7.97 | 25.17 | 4.03 |
| Linolenic acid | 13.27 | 1.031 | 4.460 |
| TSFA | 16.19 | 7.261 | 7.33 |
| TUFA | 16.19 | 7.26 | 7.33 |
| Eigenvalue | 4.56 | 1.94 | 1.26 |
| Variability (%) | 50.64 | 21.59 | 13.94 |
| Cumulative % | 50.64 | 72.23 | 86.17 |

Eigenvectors and contribution of quantitative variables in the principal component analysis.

**Fig 4.**
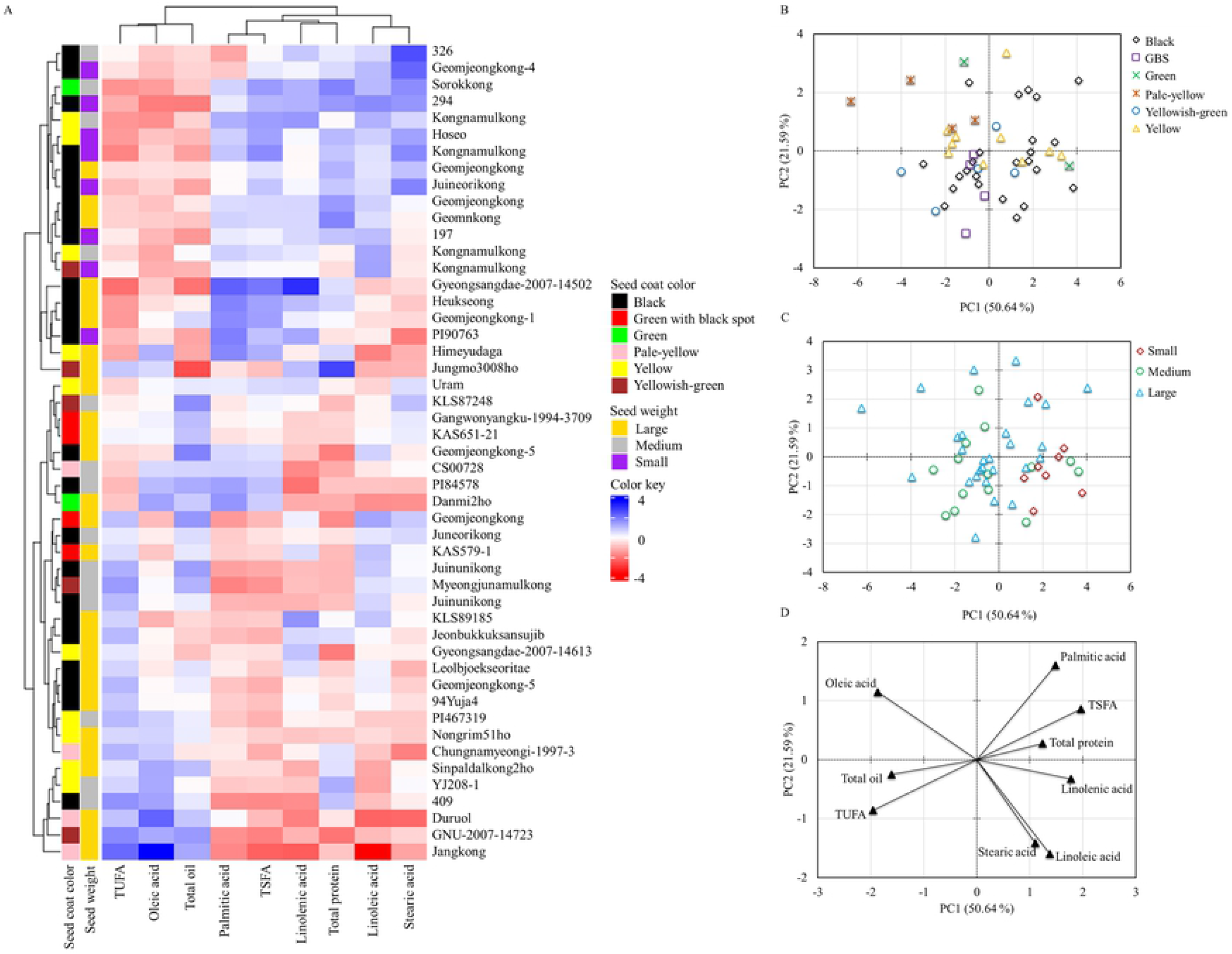
A combined heatmap and cluster analysis (A) of all the soybeans using the contents of all analyzed metabolites and score plots of observations per seed coat color (B) and seed weight (C) and loading plot of variables (D) from the principal component analysis. GBS: Green with black spot, TSFA: Total saturated fatty acid; TUFA: Total unsaturated fatty acid.

The relationship between the variables resulted from Pearson correlation analysis was also consistent with their grouping observed in the cluster analysis (Table 3). A significant but negative association (*r* = -0.714, *p* < 0.0001) was observed between total oil and total protein contents indicating soybeans with a higher protein content had a lower oil content and *vice versa*. It was depicted that the pleiotropic effects of minor and major genes associated with protein and oil contents could cause such reverse association [38]. The total oil content was also positively correlated with oleic acid (*r* = 0.444, *p* < 0.01) and negatively correlated with linolenic acid (*r* = -0.675, p < 0.0001) and palmitic acid (*r* = -0.378, *p* < 0.01). Once again, a significant but negative association (*r* = -0.936, *p* < 0.0001) was observed between oleic acid and linoleic acid. Previous studies also noted similar results and outlined that the difference in the biosynthetic pathways between these two metabolites could attribute to such trends [8, 31, 29]. Moreover, stearic acid was positively correlated with linoleic acid (*r* = 0.605) and linolenic acid (*r* = 0.254) the former association being significant (*p* < 0.0001) and the overall observation was consistent with previous findings [8, 37].

**Table 3.**

| **Variables** | **Total protein** | **Total oil** | **PA** | **SA** | **OA** | **LA** | **LLA** | **TSFA** |
| --- | --- | --- | --- | --- | --- | --- | --- | --- |
| **Total oil** | -0.714<sup>d</sup> |  |  |  |  |  |  |  |
| **Palmitic acid (PA)** | 0.236<sup>ns</sup> | -0.378<sup>b</sup> |  |  |  |  |  |  |
| **Stearic acid (SA)** | 0.249<sup>ns</sup> | -0.170<sup>ns</sup> | -0.173<sup>ns</sup> |  |  |  |  |  |
| **Oleic acid (OA)** | -0.239<sup>ns</sup> | 0.444<sup>b</sup> | -0.249<sup>ns</sup> | -0.575<sup>d</sup> |  |  |  |  |
| **Linoleic acid (LA)** | 0.088<sup>ns</sup> | -0.228<sup>ns</sup> | -0.001<sup>ns</sup> | 0.605<sup>d</sup> | -0.936<sup>d</sup> |  |  |  |
| **Linolenic acid (LLA)** | 0.392<sup>b</sup> | -0.675<sup>d</sup> | 0.368<sup>b</sup> | 0.254<sup>ns</sup> | -0.740<sup>d</sup> | 0.487<sup>c</sup> |  |  |
| **TSFA** | 0.348<sup>a</sup> | -0.446<sup>b</sup> | 0.879<sup>d</sup> | 0.316<sup>a</sup> | -0.518<sup>c</sup> | 0.291<sup>a</sup> | 0.477<sup>b</sup> |  |
| **TUFA** | -0.348<sup>a</sup> | 0.446<sup>b</sup> | -0.879<sup>d</sup> | -0.316<sup>a</sup> | 0.518<sup>c</sup> | -0.291<sup>a</sup> | -0.477<sup>b</sup> | -1.000<sup>d</sup> |

Pearson correlation coefficient (*r*) values showing the pair-wise association between quantitative variables
TSFA: Total saturated fatty acid, TUFA: Total unsaturated fatty acid.
<sup>ns</sup>Not significant, <sup>a</sup>*p* < 0.05, <sup>b</sup>*p* < 0.01, <sup>c</sup>*p* < 0.001, <sup>d</sup>*p* < 0.0001

## Conclusions

This study determined the levels of total protein, total oil and five major fatty acids in 49 soybean germplasms recently cultivated in Korea and evaluated the influences of seed coat color and seed weight on each. The results showed that neither of the factors significantly affect the total protein content. Hence, seed coat color and seed weight should not be considered as discriminant parameters during the assessment of protein content in soybeans. The total oil and individual fatty acid contents varied among the different seed coat colors and seed weights. The heatmap and PCA analysis also grouped the soybeans according to these metabolite contents and signified the importance of seed coat color and seed weight to distinguish soybean genotypes. Compared to other colored soybeans, pale-yellow soybeans were characterized by a high level of oleic acid and low levels of stearic, linoleic and linolenic acids. On the other hand, small soybeans were characterized by high levels of all individual fatty acids except oleic acid, which was significantly high in large seeds. These latter observations suggest that small and large seed soybeans could provide a wide spectrum of options to develop cultivars with modified fatty acid concentrations. Moreover, individual soybean germplasms with distinct metabolite contents identified in this study could be important sources of high-quality protein, dietary oil and fatty acids.

## Supporting information

**S1 Table**

**S2 Table**

